# Persistent Immunogenicity of Integrase Defective Lentiviral Vectors delivering membrane tethered Native-Like HIV-1 Envelope Trimers

**DOI:** 10.1101/2021.10.08.462761

**Authors:** Alessandra Gallinaro, Maria Franca Pirillo, Yoann Aldon, Serena Cecchetti, Zuleika Michelini, Antonella Tinari, Martina Borghi, Andrea Canitano, Paul F. McKay, Roberta Bona, Maria Fenicia Vescio, Felicia Grasso, Maria Blasi, Silvia Baroncelli, Gabriella Scarlatti, Celia LaBranche, David Montefiori, Mary E. Klotman, Rogier W. Sanders, Robin J. Shattock, Donatella Negri, Andrea Cara

**Affiliations:** National Center for Global Health, Istituto Superiore di Sanità, Rome, Italy; Imperial College London, Department of Infectious Disease, Norfolk Place, London, UK; Confocal Microscopy Unit NMR, Confocal Microscopy Area Core Facilities, Istituto Superiore di Sanità, Rome, Italy; Center for Gender Medicine, Istituto Superiore di Sanità, Rome, Italy; Department of Infectious Diseases, Istituto Superiore di Sanità, Rome, Italy; Department of Medicine, Division of Infectious Diseases, Duke University School of Medicine, Durham, NC, USA; Duke Human Vaccine Institute, Duke University School of Medicine, Durham, NC, USA; Viral Evolution and Transmission Unit, IRCCS Ospedale San Raffaele, 20132 Milan, Italy; Department of Surgery, Duke University School of Medicine, Durham, NC, USA; Amsterdam University Medical Centers, Amsterdam Institute for Infection and Immunity, University of Amsterdam, Amsterdam, the Netherlands; Department of Microbiology and Immunology, Weill Medical College of Cornell University, 1300 York Avenue, New York, NY, USA

## Abstract

Integrase Defective Lentiviral Vectors (IDLVs) represent an attractive vaccine platform for delivering HIV-1 antigens, given their ability to induce specific and persistent immune responses in both mice and non-human primates (NHPs). Recent advances in HIV-1 immunogen design demonstrated that native-like HIV-1 Envelope (Env) trimers that mimic the structure of virion-associated Env induce neutralization breadth in rabbits and macaques. Here, we describe the development of an IDLV-based HIV-1 vaccine expressing either soluble ConSOSL.UFO.664 or membrane-tethered ConSOSL.UFO.750 native-like Env immunogens with enhanced bNAb epitopes exposure. We show that IDLV can be pseudotyped with properly folded membrane-tethered native-like UFO.750 trimers. After a single IDLV injection in BALB/c mice, IDLV-UFO.750 induced a faster humoral kinetic as well as higher levels of anti-Env IgG compared to IDLV-UFO.664. IDLV-UFO.750 vaccinated cynomolgus macaques developed unusually long-lasting anti-Env IgG antibodies, as underlined by their remarkable half-life both after priming and boost with IDLV. After boosting with recombinant ConM SOSIP.v7 protein, two animals developed neutralization activity against the autologous tier 1B ConS virus mediated by V1/V2 and V3 glycan sites responses. By combining the possibility to display stabilized trimeric Env on the vector particles with the ability to induce sustained humoral responses, IDLVs represent an appropriate strategy for delivering rationally designed antigens to progress towards an effective HIV-1 vaccine.

## INTRODUCTION

Developing an HIV-1 vaccine that induces a durable and protective immune response remains a global health priority. The modest level of vaccine efficacy and the limited durability of antibody (Ab) responses observed in the RV144 trial highlighted the need for significant improvements^1–3^. Focused areas for improvement include the design of immunogens that could potentially drive broadly neutralizing antibodies (bNAbs) against HIV Envelope (Env)^4^ and the development of a vaccine delivery platform that can deliver those immunogens resulting in functional and durable responses.

The generation of stabilized native-like Env trimers, designed to expose bNAb epitopes while restricting the presentation of non-neutralizing epitopes, has recently contributed to major improvements in the quality of Ab responses^5,6^. These stabilized Env trimers have been shown to induce tier 2 neutralization in several animal models, including rabbits and macaques^7,8^. In order to improve the yield of well-folded native-like trimers^9,10^ and reduce the exposure of epitopes inducing non-NAbs, uncleaved pre-fusion optimized (UFO) ConSOSL.UFO antigens based on group M consensus Env sequence (Con*S*^11^) have been developed^12^. These native-like UFO trimers bear the SOS^13^ stabilizing mutation, a redesigned gp41 HR1 domain and a cleavage site linker^10^ which lead to the production of trimers with increased stability and bNAb epitopes exposure^12^. ConSOSL.UFO antigens elicited specific IgG in mice and guinea pigs and autologous tier 1B NAbs in rabbits when administered as DNA or protein, although failed at inducing heterologous neutralization^12^. In addition to the design of the immunogen, the modality of the antigen delivery plays a crucial role in the quality and durability of the induced immune responses. Several reports suggest that the long-term persistence of antigens following vaccination plays an important role in the B cell maturation process, stimulating a high degree of somatic hypermutations that results in the production of hypermutated, high affinity Abs^14,15^. Vaccination with HIV-1 Env proteins invariably leads to Ab responses that decay rapidly with a half-life of around 30-60 days^16–18^, emphasizing that considerable improvements in vaccination regimens are required to achieve durable humoral immunity against HIV-1. Self-inactivating Integrase-Defective Lentiviral Vectors (IDLVs) represent a promising vaccine platform to achieve these goals^19^. The integration deficient phenotype of IDLVs is obtained by incorporating a mutated form of the integrase gene in the recombinant parental lentiviral vector. The IDLV-encoded immunogen is therefore expressed by the unintegrated forms of vector DNA^20,21^. IDLVs have been used by us and others in several vaccine protocols to deliver viral and tumor antigens in both preclinical models^22,23^ and in cancer immunotherapeutic clinical trials^24,25^. Importantly, IDLVs provided sustained immunity against HIV-1 Env in the non-human primate (NHP) model in the absence of integration and replication^26,27^.

In addition to their potential at inducing long-term immune responses after expression of the immunogen from the unintegrated DNA, IDLVs can be harnessed to deliver immunogens through incorporation into the vector’s envelope via additional pseudotyping with heterologous viral envelope glycoproteins. In this context, we have recently shown that pseudotyping IDLV with influenza virus hemagglutinin (in addition to VSV.G) resulted in a functional Ab response in mice, with neutralization of the influenza virus persisting up to 24 weeks post-immunization^28^.

In this report, we describe the development of Simian Immunodeficiency Virus (SIV)-based IDLV delivering either soluble ConSOSL.UFO.664 (IDLV-UFO.664) or membrane-tethered ConSOSL.UFO.750 (IDLV-UFO.750) native-like HIV-1 Env trimers. Our results provide evidence that IDLV can be pseudotyped with properly folded membrane-tethered native-like Env trimers and that pseudotyped IDLV-UFO.750 induced higher IgG titers in mice compared to IDLV-UFO.664. We show that prime-boost vaccination of NHPs with IDLV-UFO.750 elicits specific and long-lived anti-Env Abs, including heterologous tier 1 NAbs and that autologous tier 1B NAbs directed to the V1/V2 and V3 glycan sites appear after boosting the IDLV-immunized animals with ConM SOSIP.v7 protein.

## RESULTS

### LV-UFO.750 produces membrane-tethered trimeric Envelopes

The basic features of the SIV-based transfer vectors used in this study are shown in **Supplementary Fig. 1**. We selected a SIV-based IDLV approach since it is more efficient than HIV-based IDLV in transducing myeloid primary cells due to the presence of viral Vpx protein in the lentiviral particles which causes degradation of the SAMHD1 restriction factor^29,30^. Truncation at amino acid 750 of the cytoplasmic tail of ConSOSL.UFO envelope and mutation of the recycling motif ^712^YSPL^715^ led to high cell surface expression levels^12^. Consistently, transfer vector pGAE-UFO.750 produced a membrane-tethered version of ConSOSL.UFO (**Fig. 1a, panels 1, 2**), which was absent in pGAE-UFO.664 transfected cells, producing a soluble version of the same Env trimer (**Fig. 1a, panels 3, 4**), as shown by Confocal Laser Scanning Microscopy (CLSM) analysis of transfected cells stained with anti-Env 2G12 bNAb.. As expected, a control plasmid expressing the parental non-engineered ConSgp160^11^, from which the ConSOSL.UFO immunogens were derived, displayed a weak plasma membrane signal (**Fig. 1a, panels 5, 6**). In contrast, all plasmids showed high and similar levels of intracellular staining (**Fig. 1a, panel 7-12**). Importantly, ConSOSL.UFO.750 showed high binding of trimer-apex quaternary-specific PGT145 or PGDM1400 bNAbs^31–33^, indicating that the Env protein is properly folded and glycosylated (**Fig. 1b**).

**Figure 1.**
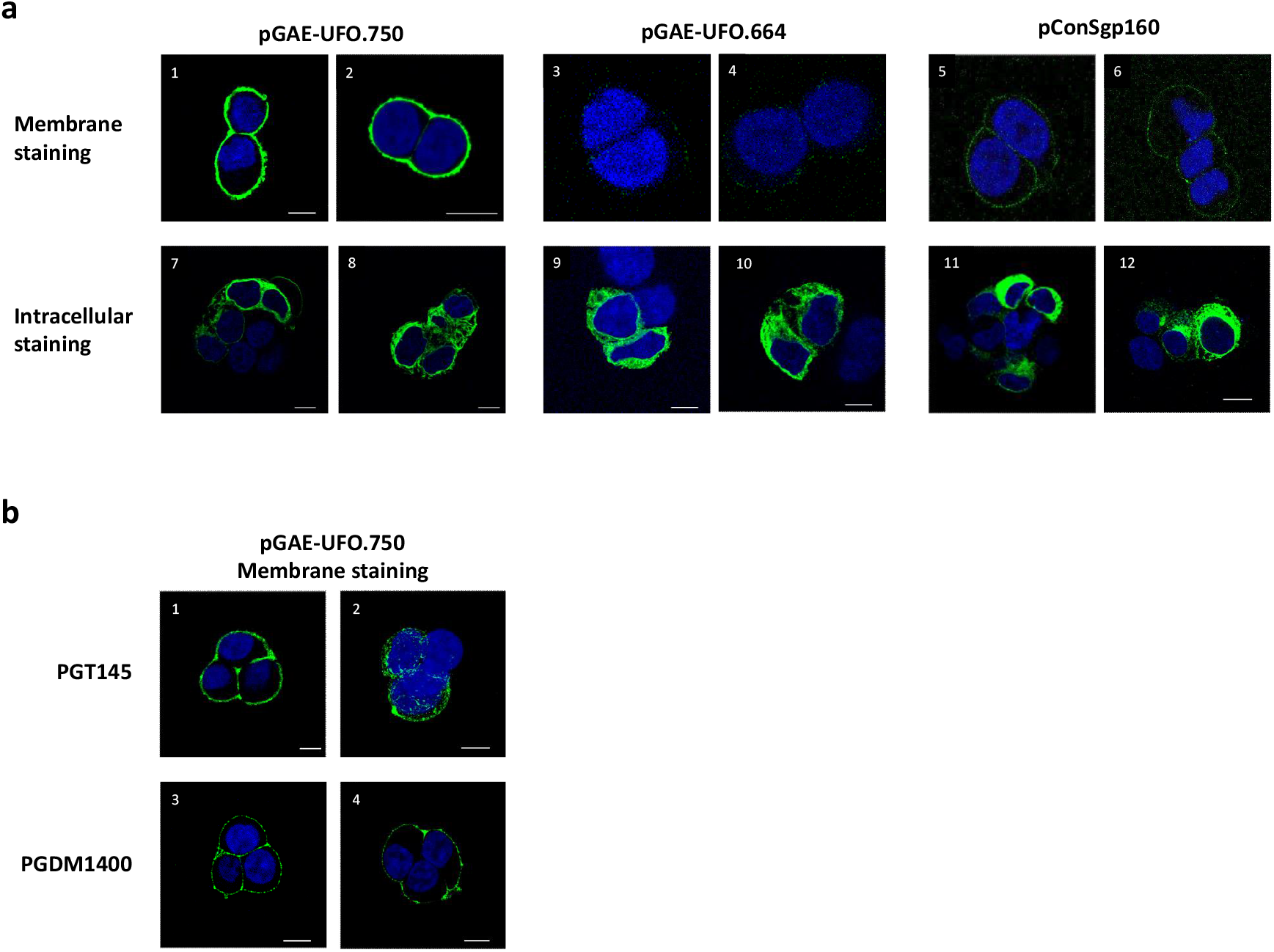
ConSOSL.UFO expression form SIV-based transfer vectors. (**a**) 293T Lenti-X cells were transfected with SIV-based transfer vectors pGAE-UFO.664 and pGAE-UFO.750 or with pConSgp160, expressing parental ConSgp160 Env. Cells were membrane (top panels) or intracellular (bottom panels) stained with **b**NAb 2G12 and analyzed by CSLM. (**b**) 293T Lenti-X cells transfected with pGAE-UFO.750 were membrane stained with trimer specific PGT145 (panels 1, 2) or PGDM1400 (panels 3, 4) bNAbs and analyzed by CSLM. Anti-human IgG Alexa Fluor 488 was used as secondary Ab. Nuclei were stained in blue by DAPI. Scale bars, 10 μm. Images represent single central optical sections. Shown are results from one representative of n = 3 experiments.

Transfer vectors were used to produce SIV-based VSV.G-pseudotyped LV-UFO.664 and LV-UFO.750, as described in the Methods section, and used to transduce 293T Lenti-X cells. Western blot analysis showed that Env was detected in the cell lysate of transduced cells using both LVs while Env was only detected in the supernatants of LV-UFO.664 transduced cells, confirming LV-UFO.664 produces soluble Env (**Fig. 2a**). We further characterized transduced cells by CLSM using anti-Env 2G12 bNAb. Both Envs were readily detected by intracellular staining (**Fig. 2b**). A weak signal was detected by extracellular staining of LV-UFO.664 transduced cells (**Fig. 2c, panels 1-6**) while LV-UFO.750 transduced cells showed strong ConSOSL.UFO.750 surface expression (**Fig. 1c, panels 7, 8**). Membrane staining with PGT145 or PGDM1400 showed that both bNAbs recognized ConSOSL.UFO.750 on the cell surface, highlighting that LV-UFO.750 produces properly folded and stable membrane-bound native-like trimers (**Fig. 1, panels 9-12**) and these data are consistent with the results obtained in cells transfected with the transfer vectors (**Fig. 1b**).

**Figure 2.**
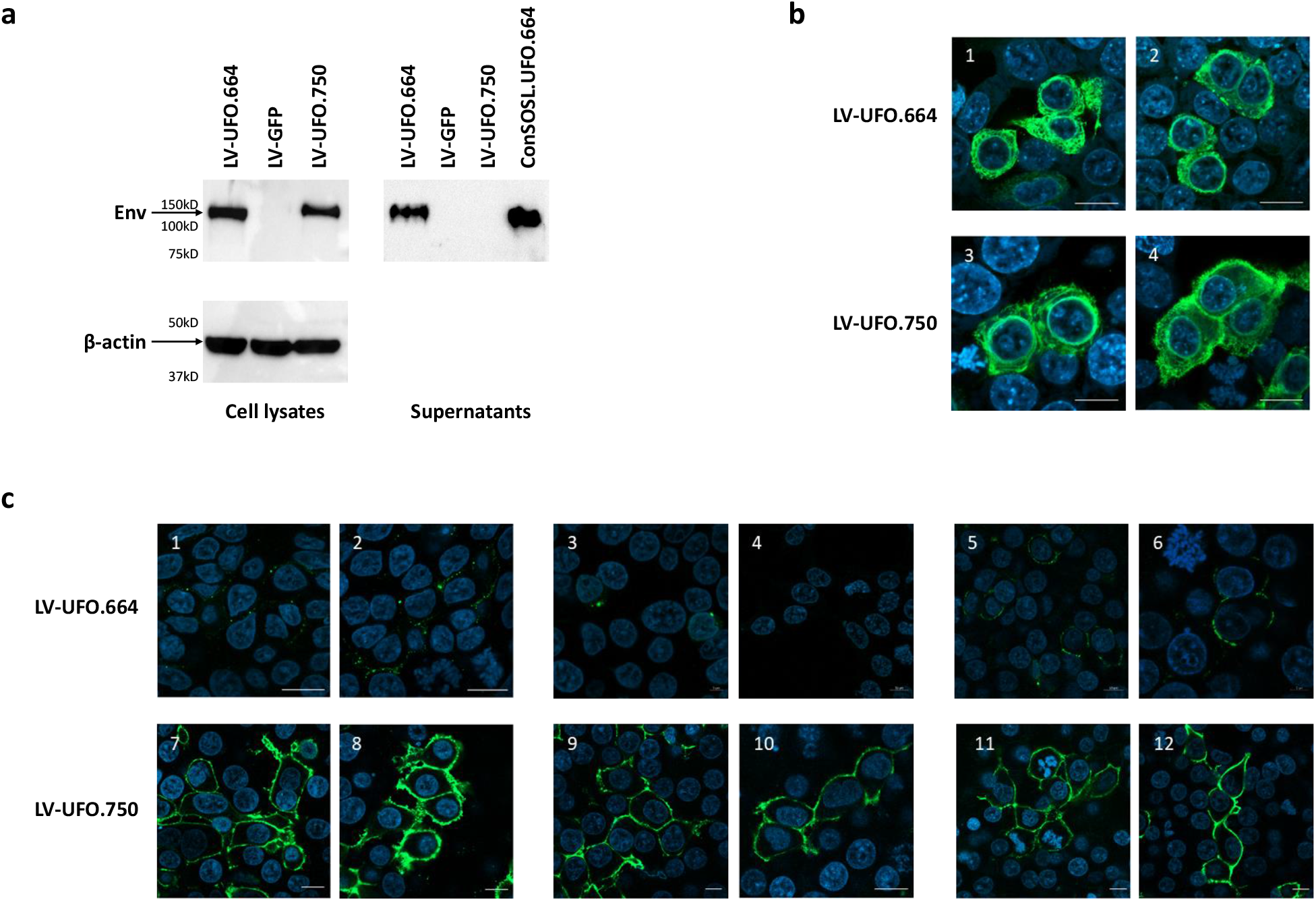
ConSOSL.UFO.664 and ConSOSL.UFO.750 expression from lentiviral vectors. 293T Lenti-X cells were transduced with 5 MOI of LV-UFO.664 or LV-UFO.750. (**a**) Six days post-transduction a WB on cells lysates (left panels) and supernatants (right panel) was performed using anti-Env 2G12 bNAb (top panels). β-actin was used as loading control (bottom panel). ConSOSL.UFO.664 recombinant protein (1 µg) and cells transduced with LV-GFP were used as positive and negative controls, respectively. (**b**) Transduced cells were also analyzed by CLSM after intracellular staining with the 2G12 bNAb. (**c**) CLSM analysis of 293T Lenti-X cells transduced with LV-UFO.664 or LV-UFO.750 after membrane staining with anti-Env 2G12 bNAb (panels 1-2, 7-8) or membrane staining with trimer-specific PGT145 (panels 3-4, 9-10) and PGDM1400 bNAbs (panels 5-6, 11-12). Anti-human IgG Alexa Fluor 488 was used as secondary Ab. Nuclei were stained in blue with DAPI and scale bars is 10 μm. Two images representing single central optical sections are shown for each NAb staining. Shown are results from one representative of n = 3 experiments.

### Native-like trimeric ConSOSL.UFO.750 pseudotypes SIV-based IDLV-UFO.750

We next produced SIV-based VSV.G-pseudotyped IDLV-UFO.664 and IDLV-UFO.750 particles for evaluating HIV-Env ConSOSL.UFO.750 incorporation. An IDLV expressing GFP (IDLV-GFP) was used as a control. Recovered and concentrated IDLV particles were normalized for p27 amount and analyzed by WB using anti-Env 2G12 and anti-Gag Abs for detection (**Fig. 3a**). While Gag protein was present in all IDLV vector preparations, Env was detected only in the IDLV-UFO.750 particles, confirming that membrane-tethered ConSOSL.UFO.750 is incorporated into IDLV-UFO.750 particles. Furthermore, truncation of Env cytoplasmic tail allowed for increased ConSOSL.UFO.750 incorporation into IDLV particles compared to parental full-length ConSgp160 (**Fig. 3b)**.

**Figure 3.**
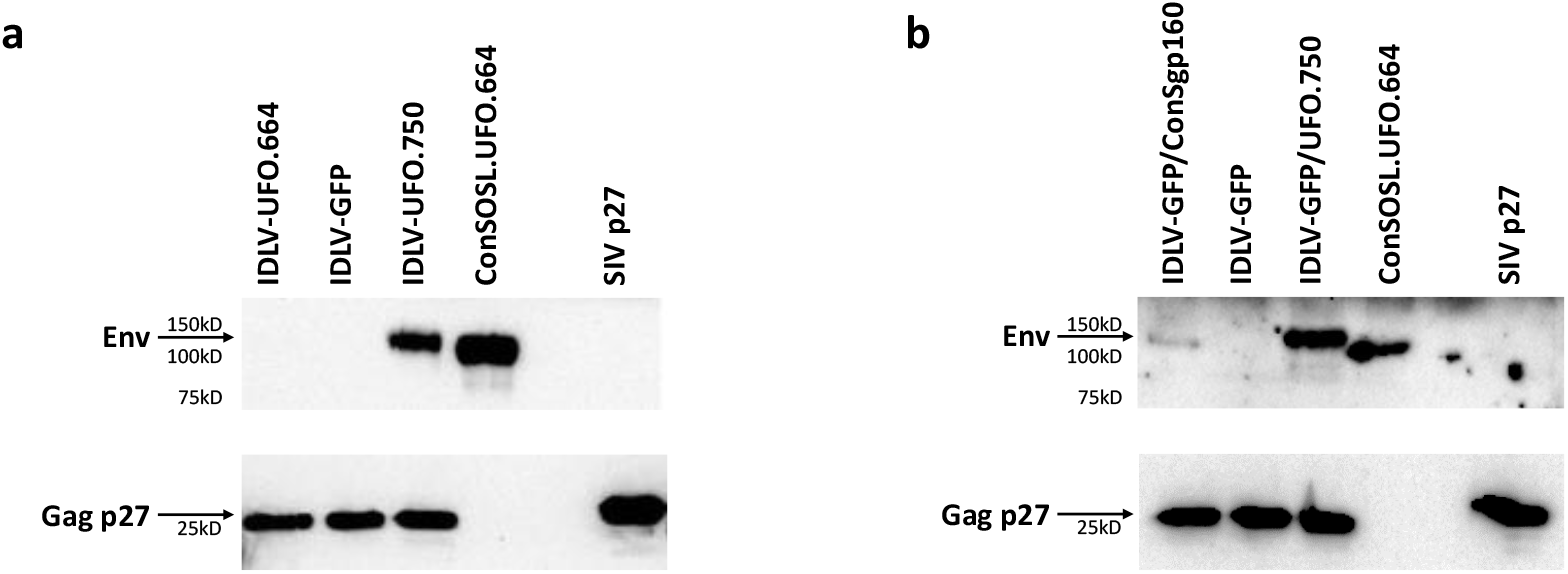
Incorporation of ConSOSL.UFO.750 envelope on IDLV particles. (**a**) Western blot of lysates from concentrated preparation of (**a**) IDLV-UFO.664 and IDLV-UFO.750 and (**b**) IDLV-GFP pseudotyped either with parental ConSgp160 or native-like ConSOSL.UFO.750 envelopes. Recombinant ConSOSL.UFO.664 and p27 SIVGag proteins (1 ug each) and IDLV-GFP were used as controls. Filters were probed with anti-Env 2G12 or anti-SIV p27 Abs. Shown are results from one representative of n = 3 experiments.

Transmission electron microscopy (TEM) observation of 293T Lenti-X cells producing IDLV-UFO.750 using 2G12 bNAb staining showed that ConSOSL.UFO.750 was found on released IDLVs and plasma membranes of producing cells (**Fig. 4a, top panels**). Importantly, we report here that the closed native-like conformation of ConSOSL.UFO.750 trimers is preserved on IDLVs surface as demonstrated by PGT145 binding observed in TEM (**Fig. 4a, bottom panels**).

**Figure 4.**
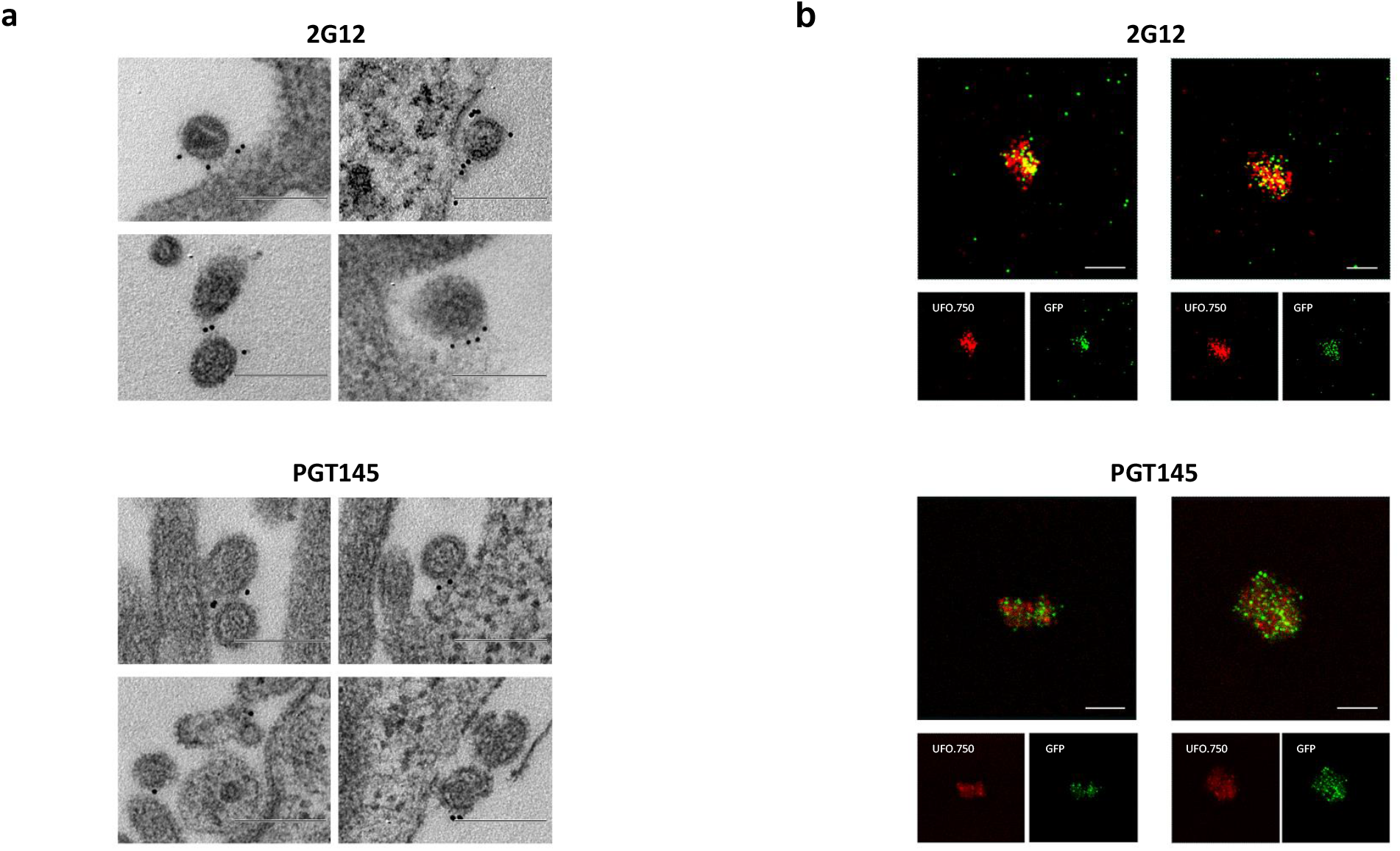
IDLV-UFO.750 is pseudotyped with native-like trimeric Envelope. (**a**) 293T Lenti-X cells producing IDLV-UFO.750 were probed with 2G12 (top panel) or PGT145 (bottom panel) bNAbs and observed by TEM. Four representative images are shown for each bNAb staining. Bars, 0.2 μm. (**b**) CLSM observation of GFP-labelled IDLV-UFO.750 obtained by adding a plasmid expressing SIV-Gag protein fused to GFP during production. Purified particles were immuno-stained with anti-Env 2G12 bNAb (top panel) or trimer-specific PGT145 bNAb (bottom panel) and anti-human IgG Alexa Fluor 647 secondary Ab (red). Top panels show the merged images in which the yellow dots represent the overlapping green (SIVGag-GFP) and red (UFO.750) signals. Bottom panels represent the individual fluorescent images. Two representative images for each Ab staining are shown. Bars, 5 μm. Shown are results from one representative of n = 2 experiments.

Presence and stability of pseudotyping ConSOSL.UFO.750 on the vector envelope surface was confirmed by CLSM using GFP-labelled SIV-based IDLV-UFO.750 (**Fig. 4b**). Labelled IDLVs were obtained by including pSIV-GagGFP plasmid^34^, in which the carboxy-terminus of SIV-Gag protein is fused to the GFP allowing for the incorporation of Gag-GFP fusion protein into IDLV-UFO.750 particles. IDLV-UFO.750 particles were stained with 2G12 or PGT145 and we observed co-localization of ConSOSL.UFO.750 and SIV-GagGFP (**Fig. 4b**). Altogether, the WB, TEM and CLSM data confirmed that IDLV-UFO.750 particles are pseudotyped with native-like ConSOSL.UFO.750 Env trimers and thus these particles may also act as virus-like particles (VLP).

Since all vectors were also pseudotyped with the VSV.G envelope glycoprotein, allowing broad tropism *in vitro* and *in vivo*^35^, we evaluated whether pseudotyping ConSOSL.UFO.750 could interfere with transduction efficiency mediated by the VSV.G. We produced GFP-expressing LV pseudotyped with ConSOSL.UFO.750 and VSV.G protein from Indiana (In) or Cocal (Co) serotypes (LV-GFP/UFO.750) or only with VSV.G (LV-GFP). 293T Lenti-X cells were transduced with escalating doses of LV-GFP/UFO.750 and LV-GFP used as control. Flow cytometry analysis showed no significant differences in LV-GFP or LV-GFP/UFO.750 transduction efficiency, regardless of the VSV.G serotype or LV dose used, indicating that pseudotyping with ConSOSL.UFO.750 did not affect the efficiency of LV transduction (**Fig. 5**).

**Figure 5.**
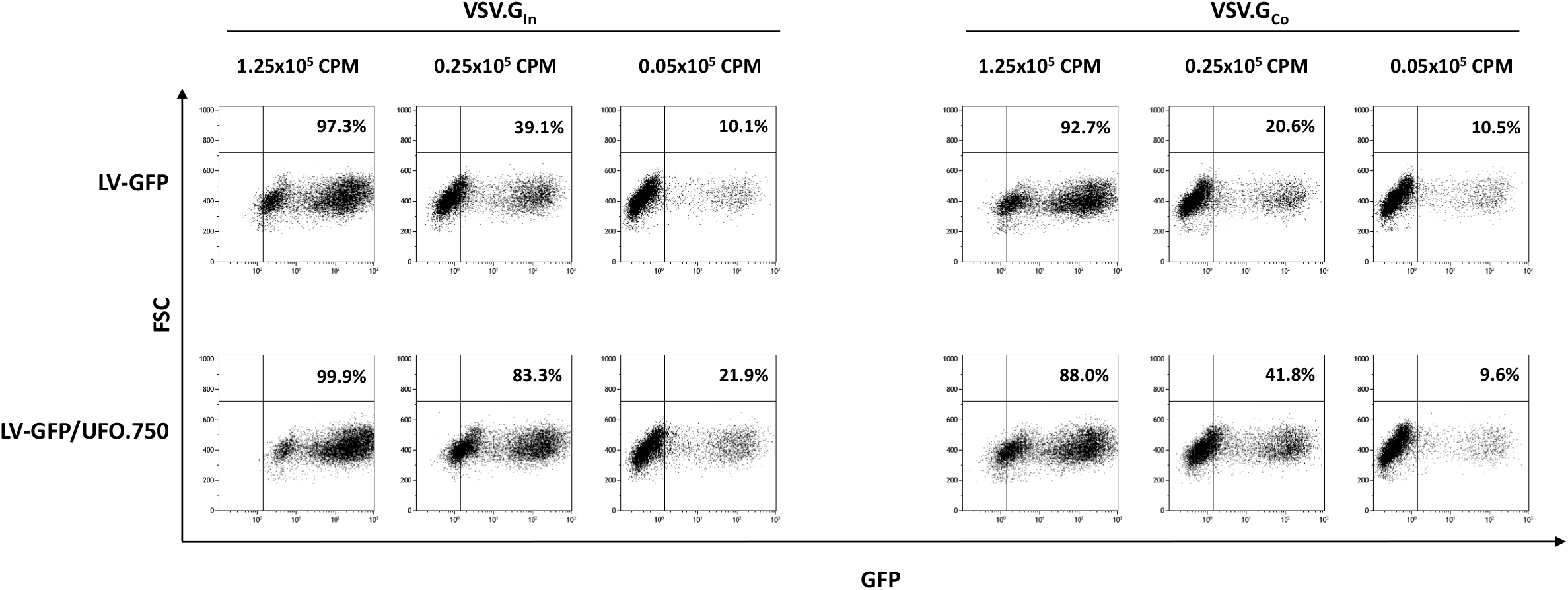
Pseudotyping ConSOSL.UFO.750 on IDLV surface does not interfere with VSV.G-mediated transduction efficiency. Top panels: Flow cytometry analysis (FSC vs GFP expression) of 293T Lenti-X cells transduced with LV-GFP pseudotyped with VSV.G serotypes from Indiana (VSV.G_In_, left panels) or Cocal (VSV.G_Co_, right panels). Bottom panels: Flow cytometry of 293T Lenti-X cells transduced with LV-GFP pseudotyped with VSV.G serotypes as above and with ConSOSL.UFO.750 (LV-GFP/UFO.750). Shown are results from one representative experiment. Percentage of positive cells is indicated. Shown are results from one representative of n = 3 experiments.

### IDLV-UFO.750 induces higher antibody titers than IDLV-UFO.664 in mice

To assess and compare the immunogenicity of IDLVs expressing pseudotyping ConSOSL.UFO.750 or soluble ConSOSL.UFO.664 Envs, we immunized BALB/c mice (n = 5 per group) once intramuscularly (i.m.) with escalating dose of either IDLV-UFO.750 or IDLV-UFO.664 (0.4×10^6^, 2×10^6^ and 10×10^6^ reverse transcriptase (RT) units/mouse). Serum anti-Env IgG were evaluated by ELISA over a 24-week period (**Fig. 6**). Results showed that a single immunization with either IDLVs elicited specific anti-Env IgG with sustained levels observed for all groups and up to the end of the study period (week 24). IDLV-UFO.750 elicited significantly higher IgG levels than IDLV-UFO.664 at 2 weeks after immunization for each dose tested. This faster humoral kinetic for the ConSOSL.UFO.750 is consistent with the previously observed DNA immunization-induced humoral response^12^. In addition, we hypothesized that the VLP characteristics of the Env pseudotyped IDLV-UFO.750 may also lead to a faster anti-Env response. These kinetic and IgG level differences were the most marked for the lowest doses of injected IDLVs. At later time point, mice vaccinated with IDLV-UFO.750 showed a higher IgG2a:IgG1 ratio compared to mice immunized with IDLV-UFO.664 (**Supplementary Fig. 2**). These data are in line with results achieved by Aldon et al.^12^ in mice immunized with plasmid DNA expressing ConSOSL.UFO.750 or ConSOSL.UFO.664 and confirm that the membrane context for Env presentation can modulate the quality of the immune response.

**Figure 6.**
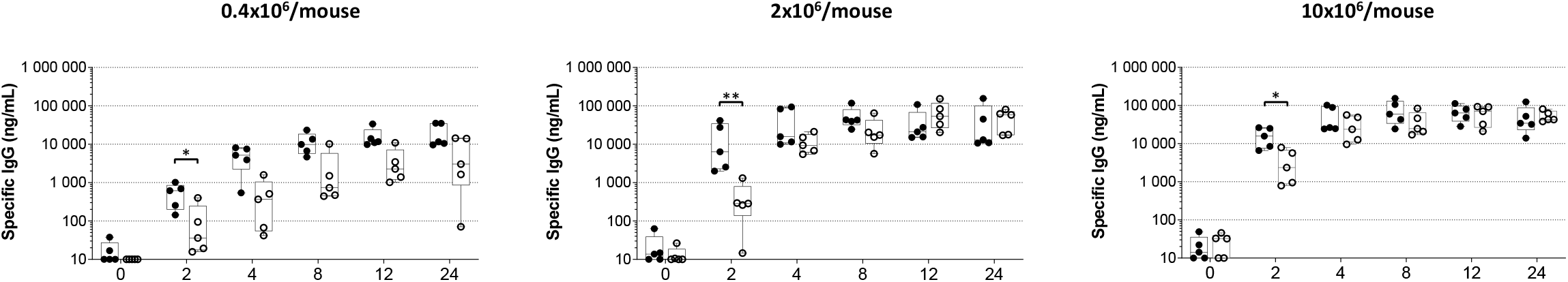
Immune response in mice immunized with IDLV delivering soluble or membrane-tethered ConSOSL.UFO. Time course of anti-ConSOSL.UFO Env IgG measured by ELISA in sera of BALB/c mice (5 mice/group) after a single immunization with escalating dose (0.4×10^6^, 2×10^6^, 10×10^6^ RT units/mouse) of either IDLV-UFO.750 (black dots) or IDLV-UFO.664 (empty dots). Samples from immunized mice were analyzed separately. Results are expressed as IgG ng/ml and shown as box and whisker plots. *p<0.05, **p<0.01, Mann-Whitney test.

### IDLV-UFO.750 induces sustained IgG responses in NHPs

Five cynomolgus macaques were vaccinated i.m. twice with IDLV-UFO.750, followed by two boosts with stabilized ConM SOSIP.v7 Env trimer^36^ adjuvanted with Monophosphoryl lipid A (MPLA). ConM SOSIP.v7 is related to ConSOSL.UFO (deriving from group M consensus sequences from 2001 and 2004, respectively^11,36^) and was chosen in the effort to improve anti-Env immune response, since previous works have shown that two administrations of native-like ConM SOSIP.v7 trimer induced anti-ConM and anti-ConS NAbs targeting the Env trimer apex in both rabbits and macaques, especially when delivered by nanoparticles^36–38^. Capture ELISA on sera from IDLV vaccinated macaques showed that priming with IDLV-UFO.750 induced specific anti-ConSOSL.UFO Env IgG in all animals (**Fig. 7**). Interestingly, following the peak at 2 weeks post-prime, the anti-Env IgG titres showed an initial contraction phase with IgG half-life of 13 weeks (between weeks 2-6 post-IDLV prime) (**Table 1**). This was followed by a long time interval showing a very low rate of decay of the vaccine-induced antibody response with IgG half-life of 58 weeks (between weeks 6-37 post-IDLV prime) (**Table 1**), similarly to what has been previously observed for vaccine-induced responses to other non-HIV vaccines and infections^39,40^. Vaccination with IDLV-UFO.750 elicited also anti-ConM Abs, albeit at much lower titers compared to anti-ConSOSL.UFO Abs. At 37 weeks after the prime, all animals were successfully boosted with IDLV-UFO.750 pseudotyped with VSV.G from non-cross-reactive Cocal serotype (VSV.G^Co^). ConSOSL.UFO-specific IgG levels peaked at 39 weeks (2 weeks post-boost) and followed a similar kinetics observed after priming, showing a higher decay by one month after the boost (between weeks 39-43, half-life of 3 weeks) and a lower decay between weeks 43-62 post-IDLV boost(half-life of 24 weeks), but reached a higher setpoint than pre-boost levels. As expected, boosting with IDLV also increased anti-ConM IgG levels which were ~1-1.5 log lower than anti-ConSOSL.UFO IgG. The first boost with ConM SOSIP.v7 induced a 1-log increase of anti-ConSOSL.UFO IgG and a 2-log increase of anti-ConM IgG. Interestingly, the anti-ConSOSL.UFO IgG response 2 weeks after the second ConM SOSIP.v7 boost was similar to that observed after the previous ConM SOSIP.v7 immunization and over time declined more rapidly than after IDLV immunizations (significant decay between weeks 64-74 and between weeks 80-94 post-ConM SOSIP.v7 protein boosts, p=0.0021 and p=0.0155, respectively) (**Table 1** and **Fig. 7**).

**Table 1:** Analysis of decay of anti-ConSOSL.UFO Env IgG response by using a piecewise linear model

|  |  |  |  |  |  |  |  |
| --- | --- | --- | --- | --- | --- | --- | --- |
| Vaccine | time interval (weeks) | Rate of decay | 95%CI<sup>a</sup> | p |  | Concentration decay/week | Half-life (weeks) |
| IDLV-UFO.750 | 2-6 | -0,053 | (-0,337; 0,231) | 0,7135 | NS | 5% | 13 |
|  | 6-37 | -0,012 | (-0,040; 0,016) | 0,4115 | NS | 1% | 58 |
| IDLV-UFO.750 | 39-43 | -0,243 | (-0,527; 0,040) | 0,0929 | NS | 2% | 3 |
|  | 43-62 | -0,029 | (-0,083; 0,025) | 0,3001 | NS | 3% | 24 |
| ConM SOSIP.v7 + MPLA | 64-74 | -0,178 | (-0,291; -0,064) | 0,0021 | ** | 16% | 4 |
| ConM SOSIP.v7 + MPLA | 76-80 | -0,313 | (-0,576; -0,051) | 0,0193 | * | 27% | 2 |
|  | 80-94 | -0,096 | (-0,174; -0,018) | 0,0155 | * | 9% | 7 |

<sup>a</sup>95%CI: 95 percent confidence intervals

**Figure 7.**
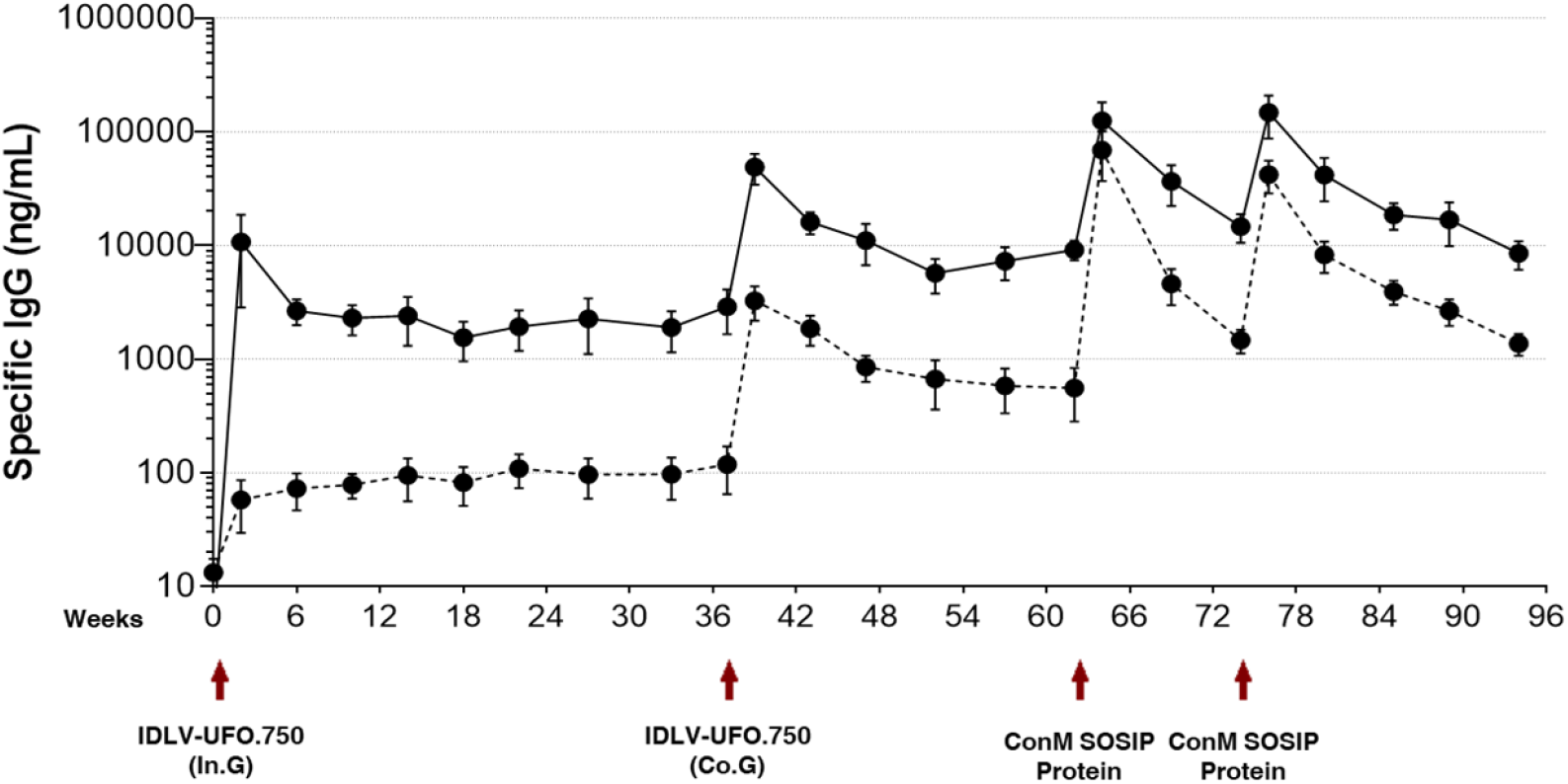
Immune response in cynomolgus macaques after prime-boost vaccination with IDLV-UFO.750 and ConM SOSIP.v7 protein. Five cynomolgus macaques were primed with IDLV-UFO.750 pseudotyped with VSV.G from Indiana serotype (In.G) and boosted 8 months after the first immunization with the same IDLV-UFO.750 pseudotyped with VSV.G from Cocal serotype (Co.G). At 62 and 74 weeks after priming, monkeys were vaccinated with ConM SOSIP.v7 protein adjuvanted with MPLA. Serum samples from vaccinated macaques were collected at the indicated time points and analyzed by capture ELISA. Results are expressed as mean concentration (ng/ml) of specific anti-ConSOSL.UFO (solid line) and anti-ConM (dotted line) IgG. Arrows indicate the immunization. Error bars indicate the standard error of the mean.

We also evaluated the presence of anti-Env IgG at mucosal sites using saliva and rectal swabs. ELISA results showed low IgG titers in the saliva in four out of five vaccinated monkeys, whereas anti-Env IgG could only be detected in one animal (AU989) in rectal samples. The kinetics of mucosal IgG was similar to that observed in the sera with peaks observed 2 weeks after each immunization (**Supplementary Table 1**).

### Prime-boost vaccination with IDLV-UFO.750 and ConM SOSIP.v7 protein induced anti-Env NAbs against V1V2 and V3 glycan site

We next tested the sera from vaccinated monkeys for the presence of vaccine-induced NAbs against a panel of HIV-1 pseudoviruses, including the tier 1A Clade C MW965.26, the tier 1B autologous ConS, and tier 2 heterologous Ce1176_A3, WITO4160.33, T-250-4, ZM233M.PB6, 6101.10, CH848.10.17, 92RW020.2, and JR-FL pseudoviruses. All monkeys developed high NAb titers against the heterologous MW965.26 virus starting from week 6 that increased after each boost (**Table 2**). We detected autologous ConS tier 1B NAb activity in 2/5 animals at week 64 (2 weeks after the first ConM SOSIP.v7 protein boost), which may suggest that priming occurred in AS304 and AU955 monkeys with the prior IDLV immunizations. For these two animals, ConS tier 1B ID50s rapidly declined to <20 but were boosted to higher levels 2 weeks after the second protein immunization (week 76) and then became undetectable by week 80. We did not detect heterologous tier 2 neutralizing activity (data not shown).

**Table 2.** Serum neutralization activity against the clade C tier 1A MW965.26 and the Consensus tier 1B ConS viruses.

| Vaccine Group | Week | MW965.26 tier 1A |  |  |  |  | ConS tier 1B |  |
| --- | --- | --- | --- | --- | --- | --- | --- | --- |
|  |  | Animal ID |  |  |  |  | Animal ID |  |
|  |  | AS304 | AT777 | AU018 | AU955 | AU989 | AS304 | AU955 |
| IDLV-UFO.750 | 0 | <20 | <20 | <20 | <20 | <20 | <20 | <20 |
|  | 2 | <20 | <20 | <20 | <20 | <20 | <20 | <20 |
|  | 6 | <20 | <20 | <20 | **29** | <20 | <20 | <20 |
|  | 10 | <20 | **27** | <20 | **104** | **90** | <20 | <20 |
|  | 22 | **47** | **71** | <20 | **203** | **378** | <20 | <20 |
| IDLV-UFO.750 | 37 | **58** | **99** | <20 | **265** | **853** | <20 | <20 |
|  | 39 | **4614** | **9278** | **930** | **35035** | **8459** | <20 | <20 |
|  | 43 | **1164** | **3076** | **418** | **7200** | **3139** | <20 | <20 |
|  | 47 | **552** | **2428** | **103** | **4013** | **1629** | <20 | <20 |
|  | 57 | **274** | **708** | **50** | **1578** | **1409** | <20 | <20 |
| ConM SOSIP.v7 + MPLA | 62 | **216** | **796** | **44** | **1159** | **1659** | <20 | <20 |
|  | 64 | **6080** | **7347** | **260** | **43740** | **7354** | **24** | **99** |
|  | 69 | **971** | **2547** | **187** | **5714** | **3589** | <20 | **24** |
| ConM SOSIP.v7 + MPLA | 74 | **467** | **1647** | **69** | **1751** | **2065** | <20 | <20 |
|  | 76 | **3618** | **4281** | **459** | **21270** | **17007** | **137** | **198** |
|  | 80 | **944** | **1936** | **309** | **7507** | **6515** | <20 | <20 |
|  | 85 | **447** | **1132** | **211** | **2413** | **1915** | <20 | <20 |
|  | 89 | **379** | **817** | **160** | **1200** | **2210** | <20 | <20 |
|  | 94 | **329** | **774** | **116** | **931** | **2917** | <20 | <20 |

Values are indicated as the serum dilution at which relative luminescence units (RLUs) were reduced 50% compared to virus control wells after subtraction of background RLUs in cell control wells. A response was considered positive if the ID50 was 3 times greater than the signal against the MLV-pseudotyped negative control virus.

Sera from monkeys AS304 and AU955 that developed autologous anti-ConS NAbs were assayed against a mini-panel of ConS mapping mutants in order to identify the epitopes targeted by these NAbs (**Table 3**). Neutralization titers were reduced by >3-fold with mutants that abrogated binding to epitopes in V1/V2 (Y173A), V2 (N156K), or V3 (N301A, G324A) regions. None of the other mutants affected the ability of tested sera to neutralize ConS pseudovirus. Thus, the mapping data suggested that the ConS-targeted neutralization activity in these animals is directed against both the V1/V2 and V3 glycan sites.

**Table 3.** Serum neutralization activity against the Consensus tier 1B ConS mutants.

| Mutant | Target | AS304 | AU955 |  |
| --- | --- | --- | --- | --- |
|  |  | week 76 | week 64 | week 76 |
| ConS | Parent | 117 | 157 | 157 |
| N276D | CD4bs | 97 | 118 | 130 |
| W680R.K683Q | MPER | 253 | 299 | 316 |
| N156K | V2 gly sens | **<30** | **<30** | **<30** |
| Y173A | V1V2 Glycan | **<30** | **<30** | **<30** |
| N295A.15 | 2G12 sens | 286 | 420 | 513 |
| N301A.10 | V3 Glycan | **32** | **53** | 125 |
| G324A | V3 Glycan | **33** | **<30** | **<30** |
| N332A.7 | V3 Glycan | 288 | 405 | 407 |
| T236K | gp120-gp41 | 103 | 150 | 172 |
| V518L | fusion peptide | 168 | 247 | 275 |
| T607S | gp120-gp41 | 78 | 95 | 108 |
| N611Q_N637Q | gp120-gp41 | 162 | 139 | 179 |

Serum samples from AS304 and AU955 monkeys were tested for the ability to neutralize a panel of ConS mutants. Values are indicated as the serum dilution at which relative luminescence units (RLUs) were reduced 50% compared to virus control wells after subtraction of background RLUs in cell control wells. A response was considered positive (shown in bold) if the ID50 was 3 times greater than the signal against the MLV-pseudotyped negative control virus.

## DISCUSSION

In this study we developed SIV-based IDLVs delivering membrane-tethered ConSOSL.UFO HIV-1 Env with the aim of inducing long-term high-magnitude NAb responses. We hypothesized that a strategy combining the sustained immunity provided by IDLV displaying native-like trimeric Env on IDLV membrane could lead to an improvement in the quality of humoral responses.

Taking advantage of the inherent features of the vector, we tested the feasibility of producing IDLV pseudotyped with membrane-tethered native-like HIV-Env trimers. As a prototype of rationally designed antigens, we selected native-like uncleaved prefusion-optimized (UFO) trimers, based on a consensus sequence of group M HIV-1 Env (ConS)^11^ and modified to produce native-like Env trimers with increased stability and exposure of epitopes recognized by bNAbs^12^. Here, we showed that IDLV particles were efficiently pseudotyped with membrane-tethered ConSOSL.UFO.750. The presence of pseudotyped HIV-Env protein on the vector was evaluated by WB and its stability and conformation by CLSM and TEM, the latter analyses confirming that the vector was pseudotyped with properly folded native-like trimers, binding to the quaternary trimer-apex-specific PGT145 bNAb. Also, incorporation of ConSOSL.UFO.750 Env protein on IDLV particles did not interfere with VSV.G pseudotyping, which is always included during vector production and essential for entry in the target cells in vitro and in vivo^41,42^.

It has been shown that muscle cells can produce Envs exposing quaternary-dependent epitopes targeted by bNAbs^12^ and act as an antigen reservoir for the induction of persistent immune responses^42^. To assess and compare the immunogenicity of membrane-tethered and soluble Envs, BALB/c mice were thus immunized i.m. with IDLV-UFO.664 and IDLV-UFO.750. Our data showed that a single i.m. immunization with the IDLV expressing membrane-tethered ConSOSL.UFO.750 induced higher IgG levels compared to IDLV expressing the soluble ConSOSL.UFO.664, which was particularly evident in the early time points and at the lowest dose of injected IDLVs. This difference in kinetics and level of IgG response between soluble and membrane-bound ConSOSL.UFO was previously observed in DNA immunization studies^12^ and indicates that the context in which the expressed transgene is presented impacts the humoral response. In addition, we showed here that IDLV-UFO.750 are pseudotyped with ConSOSL.UFO.750 and therefore may also act as VLPs. Thus, we hypothesize that this characteristic may also contribute to the earlier induction of specific Abs compared to IDLV-UFO.664 due to the presence of ConSOSL.UFO.750 on the surface of IDLVs particles which can immediately be sensed by immune cells. In contrast, ConSOSL.UFO.664 soluble protein is only expressed and released after entry and transcription of IDLV genome in the target cells.

Consequently, pseudotyped IDLV-UFO.750 was selected for downstream evaluation in NHPs. IDLV-UFO.750 induced sustained specific anti-Env IgG levels in all vaccinated cynomolgus macaques after priming. Anti-ConSOSL.UFO Env IgG titres showed a bi-phasic curve over time with an initial rapid decay in the first month after the peak followed by a second phase with long-lived anti-Env IgG responses over the next 30 weeks before the boost with half-life of 58 weeks. The long-lasting and sustained humoral response reported here is in consistent with previous studies using different HIV Envs isolates/designs and delivered by IDLV in NHPs^26,27^. In order to increase the magnitude of IgG response, animals were boosted with the same IDLV-UFO.750 pseudotyped with VSV.G_Co_ from a different serotype (Co, Cocal serotype), to avoid interference from prime-induced NAbs against VSV.G_In_ (In, Indiana serotype), as previously shown^43–45^. The IDLV-UFO.750 boost resulted in a significant increase of specific IgG levels at 2 weeks post-boost. IgG titres showed a similar bi-phasic curve than after the prime, with an initial decay after the peak followed by a second phase with durable anti-Env IgG responses over the next 20 weeks before the protein boost (half-life of 24 weeks). This is unusual, since immunization with HIV-1 Env proteins leads to IgG responses that decay rapidly with a half-life of around 30-60 days^16–18^, and suggest that long-lived plasma cells may contribute to the maintenance of immunological memory against the HIV-1 Env after IDLV-UFO.750 immunization^46^. Interestingly, this decay curve is similar to the anti-gp120 IgG decay after anti-retroviral therapy (ART) treatment to suppress HIV-1 replication in infected people, with a rapid decline followed by a more stable phase^47^, which suggests that prolonged expression at the site of injection may act as an antigen reservoir in IDLV-induced long-term immunity^42^. These results further demonstrated that IDLV prime-boost vaccination is a useful strategy to maintain sustained and high levels of antigen-specific IgG. As expected, immunization with ConM SOSIP.v7 protein boosted the IgG response although the post-peak contraction phase was significantly steeper and more rapid compared to the IDLV-induced antigen-specific IgG kinetics.

NAbs against clade C tier 1A MW965.26 virus were elicited in all vaccinated animals, with a kinetics mimicking the binding anti-ConSOSL.UFO Abs, confirming that NAb activity largely correlates the level of serum anti-Env IgG^48^. This was particularly evident after the boost with IDLV-UFO.750. Importantly, boost with ConM SOSIP.v7 induced NAbs against autologous tier 1B ConS virus in 2/5 monkeys. The induction of ConS NAbs with our immunization regimen is of interest since three administrations of DNA expressing ConSOSL.UFO.750 together with a ConSOSL.UFO.664 protein boost failed at inducing anti-ConS NAbs in rabbits^12^. Since it is very unusual to see NAb activity after the first protein administration, this data suggests that immunization with IDLV-UFO.750 acted as an effective priming. However, Brouwer et al.^37^ showed that ConM SOSIP.v7 with GLA-LSQ adjuvant could induce neutralization against ConS in one out of five rabbits after one immunization. Thus, we cannot exclude that in our protocol ConM immunization induced NAb (IgM or IgG) against ConS independently from the IDLV-UFO.750 primed B-cells. None of the immunized monkeys developed breadth against heterologous tier 2 viruses. The absence of neutralization breadth in this and other studies using ConSOSL.UFO^12^ as a vaccine suggests that further improvements in the design and selection of native-like Env immunogens are still necessary and particularly that priming immunogens such as germline targeting Envs should be used before fully glycosylated native-like trimers in order to select for rare bNAb B-cell precursors^49,50^. On the other hands, additional optimization of IDLV aiming at improving the quality of the immune response is underway. For example, increasing the native-like Env trimer density on the surface of IDLV particles has the potential to increase Env recognition by immune cells and therefore results in more potent immune responses^51,52^. Indeed, VLPs with increased density of Env are currently being pursued for vaccine development^53,54^. By displaying viral antigens on a viral surface, VLP-vaccines are strong inducers of Abs and helper T cell responses^55^. One factor limiting membrane incorporation of Env may be the presence of a long cytoplasmic tail (CT) sequence in HIV-Env (over 150 aa) compared to other viruses. In this report we have shown that truncation of native-like ConSOSL.UFO trimers at position 750 (CT of 45 aa) and mutation of known recycling motif ^712^YSPL^715^ allows Env-pseudotyping on IDLV particles. In this setting, IDLV with enhanced native-like Env incorporation represents a promising immunogen for the development of an effective and safe vaccine. Additionally, the use of the VSV.G glycoprotein in a chimeric pseudotyped VLP could also contribute to the induction of strong and prolonged immune responses thanks to the wide VSV.G tropism and to the fact that adherence of VSV.G-pseudotyped IDLV to transduced cells may lead to additional cycles of target cell transduction over time^56^. Indeed, HIV-VLP pseudotyped with VSV.G exhibited higher immunogenicity in NHP than VLP lacking VSV.G^57^.

In conclusion, we provided evidence that IDLVs, in addition to expressing the immunogen from the vector’s episomal forms, can be exploited as a VLP vaccine, enabling the exposure of stabilized Env trimers in the native-like closed conformation on the vector particles. IDLVs represent an effective candidate for delivering membrane-tethered native-like HIV-1 Env trimers and for inducing specific, sustained and functional humoral immune responses after prime-boost immunizations in NHPs.

## METHODS

### Vector Construction

SIV-based self-inactivating (SIN) transfer vector expressing GFP (pGAE-CMV-GFP-W, hereafter referred to as pGAE-GFP), Integrase-competent and Integrase-defective packaging vectors (pAdSIV3+ and pAdSIVD4V, respectively) and plasmids expressing vesicular stomatitis virus envelope glycoprotein G (VSV.G) from Indiana or Cocal serotype (phCMV-VSV.G and pMD2-Cocal.G, respectively) have been already described^44,58,59^.

ConSOSL.UFO.664 and ConSOSL.UFO.750 open reading frames (ORFs) were obtained after digestion of pCDNA3.1-ConSOSL.UFO.664 or pCDNA3.1-ConSOSL.UFO.750^12^ with BamHI/XhoI and ligated into BglII/SalI digested pGAE-GFP, removing the GFP coding sequence, to obtain pGAE-ConSOSL.UFO.664 (hereafter referred to as pGAE-UFO.664) and pGAE-ConSOSL.UFO.750 (pGAE-UFO.750), respectively. Plasmids pConSgp160 expressing ConSgp160 Env and pCDNA3-SIVGag-GFP expressing the codon optimized sequence of SIVGag fused to the GFP sequence, have been previously described^11,34^. Schematic representation of plasmids described above is shown in **Supplementary Fig. 1**.

### Lentiviral vector production

293T Lenti-X human embryonic kidney cell line (Clontech, Mountain View, CA, USA) was used for LV or IDLV production by transient transfection as previously described^22,58^. Cells were maintained in Dulbecco’s modified Eagles medium (DMEM), high glucose 4.5 g/L (Gibco, Life Technologies Italia, Monza, Italy) supplemented with 10% fetal calf serum (Corning, Mediatech, Manassas, VA, USA), 100 units/ml penicillin/streptomycin (Gibco). Briefly, 293T Lenti-X cells were seeded onto 10-cm Petri dishes and transiently transfected (CalPhos™ Mammalian Transfection Kit, Clontech Laboratories, Inc, Mountain View, CA, USA) with (i) the transfer vector plasmid expressing GFP, ConSOSL.UFO.664 or ConSOSL.UFO.750, (ii) the IN-competent or IN-defective packaging vector plasmid and (iii) the VSV.G-envelope plasmid. LV and IDLV particles were always pseudotyped with VSV.G envelope, that confers broad tropism in a wide range of cell types of many distinct host species^60^. To produce GFP-labelled IDLV-UFO.750 for confocal microscopy observation, pCDNA3-SIVGag-GFP was added to the plasmid mixture. Forty-eight hours post-transfection, the supernatant containing LVs was collected, cleared from cellular debris and passed through a 0.45 μM pore size filter (Millipore Corporation, Billerica, MA, USA). For *in vivo* animal studies, supernatants containing IDLVs were ultracentrifuged (Beckman Coulter, Fullerton, CA, USA) on a 20% sucrose cushion (Sigma Chemical, St. Louis, MO, USA) at 23,000 rpm for 2,5h at 4°C in sterile Ultra-clear centrifuge tubes (Beckman) using an SW28 swinging bucket rotor (Beckman). Pelleted vector particles were resuspended in 1X phosphate-buffered saline (PBS; Gibco) and stored at -80°C until use. Each IDLV stock was titred by the RT activity assay, and the corresponding TUs were calculated by comparing the RT activity to the one of the IDLV-GFP virions with known infectious titers, thus allowing for the determination of their infectious titer units^22,61^.

### Western blot (WB)

Pellets and supernatants of 293T Lenti-X cells transduced with LV-UFO.664 or LV-UFO.750 and IDLV concentrated preparations were resuspended in SDS loading buffer. Lysed cells, supernatants or virions were separated on 10% SDS polyacrylamide gel under reducing conditions and transferred to a nitrocellulose membrane with a Trans-Blot Turbo System (Bio-Rad, Hercules, CA, USA). Filters were saturated for 1 h with 5% nonfat dry milk in TBST (TBS with 0.1% Tween 20) and then incubated with anti-HIV-1 Env 2G12 bNAb (a gift by Dr. D. Katinger, Polymun Scientific, Klosterneuburg, Austria) or anti-HIV-1 SF2 p24 Polyclonal Ab (ARP-4250, AIDS Reagents Program, NIH) for 1 h at room temperature followed by incubation for 1h at room temperature with anti-human peroxidase conjugate IgG (Jackson ImmunoResearch, Ely, Cambridgeshire, UK) or with anti-rabbit horseradish peroxidase (HRP)-conjugated IgG (Bio-Rad, USA). Cellular pellets were also incubated with monoclonal anti-β-actin Ab (Sigma-Aldrich, St. Louis, MO, USA) and anti-mouse horseradish peroxidase (HRP)-conjugated IgG (Bio-Rad, USA). The immunocomplexes were visualized using chemiluminescence ECL detection system (WesternBright ECL, Advansta, San Jose, CA, USA). ConSOSL.UFO.664 protein (a gift by Dr. D. Katinger, Polymun Scientific) and SIVmac239 p27 Recombinant Protein (cat # 13446, AIDS Reagents Program, NIH) were used as positive controls, whereas cells transduced with LV-GFP or concentrated preparations of IDLV-GFP were used as negative controls.

### Confocal laser scanner microscopy (CLSM)

293T Lenti-X cells (2.5×10^4^/well) were seeded in 24-well microplates onto 12-mm cover glasses previously treated with L-polylysine (Sigma) and transiently transfected with pGAE-UFO.664, pGAE-UFO.750 or pConSgp160 using the CalPhos™ Mammalian Transfection Kit (Clontech Laboratories) or transduced with 5 MOI of LV-UFO.664 or LV-UFO.750.. Twenty-four hours after the transfection and forty-height hours post-transduction, cells were washed and directly stained, prior to fixation, with 2G12, PGT145 or PGDM1400 bNAbs followed by AlexaFluor 488 Goat anti-human IgG (Jackson ImmunoResearch) to detect the membrane expression, or fixed with paraformaldehyde 3%, permeabilized with Triton 0.5% and stained with the above-mentioned primary and secondary Abs. The coverslips were mounted with Vectashield antifade mounting medium-containing DAPI (Vector Labs, Burlingame, CA, USA) on the microscope slides.

To detect pseudotyping ConSOSL.UFO.750 on IDLV particles, 3×10^4^ TU of GFP-labelled IDLV-UFO.750 were exposed to clean glass coverslips previously treated with 10 ug/ml Polybrene (Millipore, Burlington, MA, USA). Coverslips were rinsed and directly immunostained with anti-Env 2G12 or PGT145 bNAbs and AlexaFluor 647 Goat anti-human IgG secondary Ab (Jackson ImmunoResearch) prior to fixation. CLSM observations were performed on a Leica TCS SP2 AOBS apparatus (Leica Microsystems, Wetzlar, Germany) equipped with a 63x/1,3 NA using excitation spectral laser lines at 405, 488 and 633, and using the confocal software (Leica, Wetzlar, Germany) (**Fig. 1, and Fig. 4b)** or with a Zeiss LSM980 apparatus (Zeiss, Oberkochen, Germany), equipped with a 63x/1,4 WD 0,17mm oil objective, Airyscan2 and excitation spectral laser lines at 405, 488, 633 nm (**Fig. 2c and 2b**). Image acquisition and processing was carried out using Zen Blue edition 3.3 (Zeiss) and Adobe Photoshop CS5 software programs (Adobe Systems, San Jose, CA, USA). Cells stained only with the fluorochrome-conjugated secondary antibody were used to set up acquisition parameters. Signals from different fluorescent probes were taken in sequential scanning mode. Several fields of view (>200 cells) were analyzed for each labeling condition, and representative results are shown.

### Transmission electron microscopy (TEM) analysis

293T Lenti-X cells (3×10^5^/well) were seeded in 6 well plate and transfected to produce IDLV-UFO.750 as described above. Forty-eight hours after transfection, confluent monolayers of 293T Lenti-X cells were stained with 2G12 or PGT145 bNAbs and Goat Anti-Human IgG H&L (10 nm Gold) secondary Ab (Abcam, Cambridge, UK) and then fixed in 2.5% glutaraldehyde in cacodylate buffer 0.1 M, pH 7.2, washed and post-fixed in 1% OsO4 in the same buffer. After washing, fixed specimens were dehydrated through a graded series of ethanol solutions and embedded in Agar 100 resin (Agar Scientific, Essex, UK). Ultrathin sections were collected on 200-mesh copper grids and counterstained with uranyl acetate and lead citrate. Sections were observed with a Philips 208S transmission electron microscope at 100 kV.

### Flow Cytometry

293T Lenti-X cells were transduced with escalating doses (0.05×10^5^ - 0.25×10^5^ - 1.25×10^5^ RT units) of SIV-based LV-GFP pseudotyped with VSV.G from Indiana (In.G) or Cocal (Co.G) serotype and with ConSOSL.UFO.750. IDLV-GFP pseudotyped only with either VSV.G were used as control. Transduced cells were analyzed by flow cytometry to evaluate GFP expression. GFP expression was evaluated by measuring fluorescence using the FACSCalibur (BD Biosciences, Milan, Italy), and data were analyzed using Kaluza Analysis Software (Beckman).

### Mouse immunization protocol

BALB/c mice were purchased from Charles River (Charles River, Calco, Como, Italy) and housed under specific pathogen-free conditions in the animal facility of the Istituto Superiore di Sanità (ISS, Rome, Italy). All animal procedures have been performed in accordance with European Union guidelines and Italian legislation for animal care. All animal studies were authorized by the Italian Ministry of Healthy and reviewed by the Service for Animal Welfare at ISS (Authorization n. 314/2015-PR of 30/04/2015). Groups of five mice were immunized once intramuscularly (i.m.) with escalating doses (0.4×10^6^ - 2×10^6^ - 10×10^6^ RT units/mice) of SIV-based IDLV-UFO.664 or IDLV-UFO.750. Naïve mice were used as negative controls. Blood retro orbital sampling was performed prior to immunization, two weeks post-immunization and at monthly intervals with glass Pasteur pipettes and sera were tested for the presence of anti-UFO antibodies by ELISA assay (described below).

### Non-human primate (NHP) immunization protocol

Five adult male cynomolgus monkeys (*Macaca fascicularis*) were housed under specific pathogen-free conditions in the animal facility of the Istituto Superiore di Sanità (ISS, Rome, Italy) according to the European Union guidelines and Italian legislation for non-human primate care. All animal studies were authorized by the Italian Ministry of Healthy and reviewed by the Service for Animal Welfare at ISS (Authorization n. 305/2016-PR of 24/03/2016). Macaques were immunized i.m. with 3×10^8^ TU/animal of IDLV-UFO.750 in 1 ml injection volume divided into two sites (left and right thighs). Thirty-seven weeks after the prime, animals were boosted with the same IDLV-UFO.750 vector but pseudotyped with a different VSV.G serotype (VSV.G Indiana for the prime, VSV.G Cocal for the boost). Six months after the second IDLV administration, monkeys received 45 µg of ConM SOSIP.v7 protein (a gift by Dr. D. Katinger, Polymun Scientific) adjuvanted with 650 µg of MPLA (a gift by Dr. D. Katinger, Polymun Scientific), followed three months later by a fourth vaccination with the same adjuvanted protein. Peripheral blood cells, sera, plasma, saliva, and rectal swabs were obtained prior to immunization, at two weeks after each immunization and every four weeks thereafter after sedation with ketamine hydrochloride (10 mg/kg) for routine hematological, biochemical and immunological determinations. All experimental procedures on macaques, were performed under direct supervision of the veterinarian staff, in accordance with the institutional policies for animal health and wellbeing.

Mucosal Abs were eluted from the Weck-Cel sponges (Beaver-Visitec, Waltham, MA, USA) with the antibody extraction buffer prepared by adding the protease inhibitor cocktail set I (Millipore) to 2x PBS. Sponges were placed into the top chamber of a Spin-X® column (Costar, Washington, USA) and incubated at RT for 10 minutes. The Spin-X® column was then centrifuged at 12,000 g for 15 min to remove large debris and isolate the fluid containing the high salt eluted antibody which was then frozen at −80°C.

### Measurement of binding antibodies by ELISA

To detect anti-Env binding Abs in mice vaccinated with either IDLV-UFO.664 or IDLV-UFO.750, ninety-six well Maxisorp plates (Nalge Nunc, Rochester, NY, USA) were coated with ConSOSL.UFO.664 recombinant protein (1 µg/ml in PBS 1X) overnight at 4°C. After washing (PBS 1X, 0,05% Tween-20) and blocking with assay buffer (PBS 1X, 1% BSA, 0.05% Tween-20), serial dilutions (1:100, 1:1000; 1:10000) of serum from individual mice were added to wells in triplicate and incubated for 1 h at 37°C. The plates were washed and HRP-conjugated goat anti-mouse IgG (Southern Biotech, Birmingham, AL, USA) was added to the wells for 1 h at 37°C. Then, SureBlue TMB Peroxidase solution (KPL, Gaithersburg, MD, USA) was added for 5 minutes at room temperature, followed by 50 µl/well of TMB stop solution (KPL). Five-fold dilutions of mouse IgG (Southern Biotech) were used to develop standard curves.

For NHP samples, capture ELISA was performed. Maxisorp plates (Nalge Nunc) were coated with 2.5 µg/ml of mouse anti-human c-Myc 9E10 mAb (produced in house) or goat anti-human IgG (1:2000, Southern Biotech) for the standard. After overnight at 4°C, plates were washed and blocked with casein buffer (Thermo Scientific, Waltham, MA, USA) at 37°C for 1h. After washing, 1 µg/ml of ConSOSL.UFO.664 MycHis Tagged protein (produced in house) was incubated for 1h at 37°C. Three five-fold dilutions of Cynomolgus IgG standard (Molecular Innovations, Novi, MI, USA) were added in triplicate and incubated for 1h at 37°C. Then, biotinylated mouse anti-monkey IgG (Southern Biotech) was added 1:50000. After 1h at 37°C, Poly-HRP 40 (1:1000, Fitzgerald, Acton, MA, USA) was added for 45 minutes at 37°C and detection was performed with SureBlue TMB Peroxidase solution (KPL) for 5 minutes at room temperature, followed by 50 µl/well of TMB stop solution (KPL). The concentration of both mouse and simian IgG antibodies was calculated relative to the standard using a 5-parameter fit curve (Softmax, Molecular Devices, Sn Jose, CA, USA) and results are expressed as µg/mL of HIV-specific antibodies (IgG) and for each group of immunization.

### HIV neutralization assays

Neutralization of Env-pseudotyped viruses was measured in 96-well culture plates using Tat-regulated firefly luciferase (Luc) reporter gene expression to quantify reductions of virus infection in TZM-bl cells^62,63^. Serum samples were heat inactivated (1 hour at 56°C) and assayed at four four-fold dilutions starting with 1:10. Each serum were incubated with 1.5×10^5^ relative luminescence units (RLU) of HIV MW965 (tier 1A), ConS (autologous, tier 1B), and heterologous HIV Ce1176_A3.LucR.T2A.ecto, WITO4160.33, T-250-4, ZM233M.PB6, 6101.10, CH848.10.17, 92RW020.2, and JR-FL pseudoviruses. Sera that showed anti-ConS tier 1B neutralization titer of about 100 were tested against a panel of 11 ConS mutants (N156K, Y173A, N301A.10, G324A, N332A.7, N295A.15, N276D, V518L, T236K, T607S, N611Q, N637Q) to evaluate the neutralization profile. Neutralizing activity against MLV-pseudotyped virus was also tested as a negative control for non-HIV-specific inhibitory activity in the assays (data not shown). Luciferase activity was measured using Bright-Glo reagent (Promega Corporation, Madison, WI, USA). Neutralizing antibody titers were expressed as the serum dilution at which relative luminescence units (RLUs) were reduced 50% compared to virus control wells after subtraction of background RLUs in cell control wells. A response was considered positive if the post immunization ID50 was 3 times higher than the pre-immune ID50.

### Statistical Analysis

The comparison of antibody response in the immunized mice was assessed by the nonparametric Mann-Whitney test, using GraphPad Prism v9.1.2 (GraphPad Software, San Diego, CA, USA). Statistical tests were conducted two-sided at an overall significance level of p = 0.05. Antibodies concentrations in the immunized monkeys were log transformed. A piecewise linear model was carried out within a structural equation modelling framework to estimate the rate of decay of the antibodies at different time intervals^64^. To account for the intra-individual correlation a latent variable, at the animal level, was incorporated in the model. The half-life was estimated as follows: 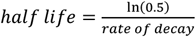. All analyses were carried out in Stata 16.1^65^ (StataCorp. 2019. Stata: Release 16. Statistical Software. College Station, TX: StataCorp LLC).

## ACKNOWLEDGEMENTS

We are grateful to Dr. Omar Leoni and Dr. Gianluca Panzini for veterinary assistance and to all personnel responsible for the animal facility: Antonio Di Virgilio, Luigia Cancemi, Maurizio Chiodi, Anna Rita Lamagna, Isabella Marcucci, and Francesca Ubaldi. We are grateful to Stefania Donnini for secretarial assistance; Ferdinando Costa and Patrizia Cocco for technical support. We also thank Iole Farina for her help in NHP samples processing. We thank Dr. Dietmar Katinger for providing recombinant human monoclonal antibody to HIV-1 gp120, 2G12 (#AB002); recombinant human monoclonal antibody to HIV-1 gp120, PT145; ConM SOSIP.v7 and ConSOSL.UFO.664 Protein and MPLA adjuvant. We thank Marit J. van Gils and Emma I. M. M. Reiss for donating recombinant human monoclonal antibody to HIV-1 gp120, PGDM1400. This work was supported in part by grants from the National Institute of Allergy and Infectious Diseases (NIAID; 1P01AI110485-01A1 to M.E.K.) and Italian Ministry of Health Ricerca Finalizzata (PE-2011-02347035 to A. Cara and PE-2016-02364927). This project has received funding from the European Union’s Horizon 2020 research and innovation program under grant agreement no. 681137 (EAVI2020) and the European Union’s Seventh Programme for Research, Technological Development and Demonstration under grant agreement no. 280873 (ADITEC; to A. Cara and D.N.). We thank Fondation Dormeur, Vaduz for the donation of laboratory instruments relevant to this project to the Istituto Superiore di Sanità. The following reagents were obtained through the NIH HIV Reagent Program, Division of AIDS, NIAID, NIH: Polyclonal Anti-Human Immunodeficiency Virus Type 1 SF2 p24 (antiserum, Rabbit), ARP-4250, contributed by DAIDS/NIAID; produced by BioMelecular Technologies; SIVmac239 p27 Recombinant Protein from NIAID, DAIDS (cat# 13446).

## AUTHOR CONTRIBUTIONS

A.G. participated in study design, performed the majority of the experiments, analyzed the data, and wrote the manuscript. M.F.P. and R.B. contributed to vector design, construction, and preparation. Y.A. performed ELISA assay and contributed to data analysis. P.F.M. performed ELISA assay. S.C. performed the CLSM experiments. Z.M. and A.T. performed TEM experiments. M. Borghi performed the RT assay and contributed to mice immunization and NHPs samples processing. A. Canitano and F.G. performed WB experiments. S.B. coordinated NHPs immunization and samples processing. C.L. and D.M. designed and analyzed the neutralization and epitope mapping assays. R.J.S. contributed to study design and data analysis. M.B., G.S., M.E.K. and R.W.S. participated in study design. D.N. contributed to study design, data analysis and interpretation of the data, and editing of the manuscript. A. Cara oversaw the planning and direction of the project, including analysis and interpretation of the data and editing of the manuscript.

## ADDITIONAL INFORMATIONS

**Supplementary information** is available for this paper at

**Correspondence** and requests for materials should be addressed to A. Cara.

## DATA AVAILABILITY

The data that support the findings of this study are available from the corresponding author upon reasonable request.

## REFERENCES

1. Rerks-Ngarm, S. et al. Vaccination with ALVAC and AIDSVAX to Prevent HIV-1 Infection in Thailand. N. Engl. J. Med. 361, 2209–2220 (2009).

2. Haynes, B. F. . et al. Immune-Correlates Analysis of an HIV-1 Vaccine Efficacy Trial. N. Engl. J. Med. 366, 1275–1286 (2012).

3. Gray, G. E. et al. Vaccine Efficacy of ALVAC-HIV and Bivalent Subtype C gp120–MF59 in Adults. N. Engl. J. Med. 384, 1089–1100 (2021).

4. Haynes, B. F. & Burton, D. R. Developing an HIV vaccine. Science 355, 1129–1130 (2017).

5. Sanders, R. W. et al. A Next-Generation Cleaved, Soluble HIV-1 Env Trimer, BG505 SOSIP.664 gp140, Expresses Multiple Epitopes for Broadly Neutralizing but Not Non-Neutralizing Antibodies. PLoS Pathog. 9, (2013).

6. Sanders, R. W. et al. HIV-1 neutralizing antibodies induced by native-like envelope trimers. Science 349, (2015).

7. Sanders, R. W. & Moore, J. P. Native-like Env trimers as a platform for HIV-1 vaccine design. Immunol. Rev. 275, 161–182 (2017).

8. Saunders, K. O. et al. Vaccine Induction of Heterologous Tier 2 HIV-1 Neutralizing Antibodies in Animal Models. Cell Rep. 21, 3681–3690 (2017).

9. Kovacs, J. M. et al. Stable, uncleaved HIV-1 envelope glycoprotein gp140 forms a tightly folded trimer with a native-like structure. Proc. Natl. Acad. Sci. U. S. A. 111, 18542–18547 (2014).

10. Kong, L. et al. Uncleaved prefusion-optimized gp140 trimers derived from analysis of HIV-1 envelope metastability. Nat. Commun. 7, 1–15 (2016).

11. Liao, H. et al. A Group M Consensus Envelope Glycoprotein Induces Antibodies That Neutralize Subsets of Subtype B and C HIV-1 Primary Viruses. Virology 353, 268–282 (2006).

12. Aldon, Y. et al. Rational Design of DNA-Expressed Stabilized Native-Like HIV-1 Envelope Trimers. Cell Rep. 24, 3324–3338 (2018).

13. Sanders, R. W. et al. Stabilization of the Soluble, Cleaved, Trimeric Form of the Envelope Glycoprotein Complex of Human Immunodeficiency Virus Type 1. J. Virology 76, 8875–8889 (2002).

14. Fouda, G. G. A. et al. Mucosal Immunization of Lactating Female Rhesus Monkeys with a Transmitted/Founder HIV-1 Envelope Induces Strong Env-Specific IgA Antibody Responses in Breast Milk. J. Virol. 87, 6986–6999 (2013).

15. Baumjohann, D. et al. Persistent Antigen and Germinal Center B Cells Sustain T Follicular Helper Cell Responses and Phenotype. Immunity 38, 596–605 (2013).

16. Anderson, K. P. et al. Effect of dose and immunization schedule on immune response of baboons to recombinant glycoprotein 120 of HIV-1. J. Infect. Dis. 160, 960–969 (1989).

17. Gilbert, P. B. et al. Correlation between immunologic responses to a recombinant glycoprotein 120 vaccine and incidence of HIV-1 infection in a phase 3 HIV-1 preventive vaccine trial. J. Infect. Dis. 191, 666–677 (2005).

18. Klasse, P. J., Sanders, R. W., Cerutti, A. & Moore, J. P. How can HIV-type-1-Env immunogenicity be improved to facilitate antibody-based vaccine development? AIDS Res. Hum. Retroviruses 28, 1–15 (2012).

19. Negri, D. R. M., Michelini, Z. & Cara, A. Toward integrase defective lentiviral vectors for genetic immunization. Curr. HIV Res. 8, 274–281 (2010).

20. Vargas, J., Gusella, G. L., Najfeld, V., Klotman, M. E. & Cara, A. Novel integrase-defective lentiviral episomal vectors for gene transfer. Hum. Gene Ther. 15, 361–372 (2004).

21. Wanisch, K. & Yáñez-Muñoz, R. J. Integration-deficient lentiviral vectors: A slow coming of age. Mol. Ther. 17, 1316–1332 (2009).

22. Negri, D. R. M. et al. Successful immunization with a single injection of non-integrating lentiviral vector. Mol. Ther. 15, 1716–1723 (2007).

23. Coutant, F., Frenkiel, M. P., Despres, P. & Charneau, P. Protective antiviral immunity conferred by a nonintegrative lentiviral vector-based vaccine. PLoS One 3, (2008).

24. Somaiah, N. et al. First-in-class, first-in-human study evaluating LV305, a dendritic-cell tropic lentiviral vector, in sarcoma and other solid tumors expressing NY-ESO-1. Clin. Cancer Res. 25, 5808–5817 (2019).

25. Pollack, S. M. et al. First-in-Human Treatment with a Dendritic Cell-Targeting Lentiviral Vector-expressing NY-ESO-1, LV305, Induces Deep, Durable Response in Refractory Metastatic Synovial Sarcoma Patient. J. Immunother. 40, 302–306 (2017).

26. Negri, D. et al. Immunization with an SIV-based IDLV expressing HIV-1 Env 1086 Clade C elicits durable humoral and cellular responses in rhesus macaques. Mol. Ther. 24, 2021–2032 (2016).

27. Blasi, M. et al. Immunogenicity, safety, and efficacy of sequential immunizations with an SIV-based IDLV expressing CH505 Envs. npj Vaccines 5, 1–12 (2020).

28. Gallinaro, A. et al. Integrase defective lentiviral vector as a vaccine platform for delivering influenza antigens. Front. Immunol. 9, (2018).

29. Laguette, N. et al. SAMHD1 is the dendritic – and myeloid – cell – specific HIV – 1 restriction factor counteracted by Vpx. Nature 474, 654–657 (2011).

30. Negri, D. R. M. et al. Simian immunodeficiency virus-Vpx for improving integrase defective lentiviral vector-based vaccines. Retrovirology 9 (2012).

31. Yasmeen, A. et al. Differential binding of neutralizing and non-neutralizing antibodies to native-like soluble HIV-1 Env trimers, uncleaved Env proteins, and monomeric subunits. Retrovirology 11, 1–17 (2014).

32. Walker, L. M. et al. Broad neutralization coverage of HIV by multiple highly potent antibodies. Nature 477, 466–470 (2011).

33. Sok, D. et al. Recombinant HIV envelope trimer selects for quaternary-dependent antibodies targeting the trimer apex. Proc. Natl. Acad. Sci. U. S. A. 111, 17624–17629 (2014).

34. Gallinaro, A. et al. Development and Preclinical Evaluation of an Integrase Defective Lentiviral Vector Vaccine Expressing the HIVACAT T Cell Immunogen in Mice. Mol. Ther. - Methods Clin. Dev. 17, 418–428 (2020).

35. Naldini, L. et al. In Vivo Gene Delivery and Stable Transduction of Nondividing Cells by a Lentiviral Vector. Science 272, 263–267 (1996).

36. Sliepen, K. et al. Structure and immunogenicity of a stabilized HIV-1 envelope trimer based on a group-M consensus sequence. Nat. Commun. 10, (2019).

37. Brouwer, P. J. M. et al. Enhancing and shaping the immunogenicity of native-like HIV-1 envelope trimers with a two-component protein nanoparticle. Nat. Commun. 10, (2019).

38. Antanasijevic, A. et al. Structural and functional evaluation of de novo-designed, two-component nanoparticle carriers for HIV Env trimer immunogens. PLoS Pathogens 16 (2020).

39. Dalby, T., Petersen, J. W., Harboe, Z. B. & Krogfelt, K. A. Antibody responses to pertussis toxin display different kinetics after clinical Bordetella pertussis infection than after vaccination with an acellular pertussis vaccine. J. Med. Microbiol. 59, 1029–1036 (2010).

40. White, M. T. et al. A combined analysis of immunogenicity, antibody kinetics and vaccine efficacy from phase 2 trials of the RTS,S malaria vaccine. BMC Med. 12, (2014).

41. Negri, D. R. M. et al. Transduction of human antigen-presenting cells with integrase-defective lentiviral vector enables functional expansion of primed antigen-specific CD8+ T cells. Hum. Gene Ther. 21, 1029–1034 (2010).

42. Lin, Y. Y. et al. Skeletal Muscle Is an Antigen Reservoir in Integrase-Defective Lentiviral Vector-Induced Long-Term Immunity. Mol. Ther. - Methods Clin. Dev. 17, 532–544 (2020).

43. Rose, N. F., Roberts, A., Buonocore, L. & Rose, J. K. Glycoprotein Exchange Vectors Based on Vesicular Stomatitis Virus Allow Effective Boosting and Generation of Neutralizing Antibodies to a Primary Isolate of Human Immunodeficiency Virus Type 1. J. Virol. 74, 10903–10910 (2000).

44. Beignon, A.S. et al. Lentiviral Vector-Based Prime/Boost Vaccination against AIDS: Pilot Study Shows Protection against Simian Immunodeficiency Virus SIVmac251 Challenge in Macaques. J. Virol. 83, 10963–10974 (2009).

45. Blasi, M. et al. IDLV-HIV-1 Env vaccination in non-human primates induces affinity maturation of antigen-specific memory B cells. Commun. Biol. 1, (2018).

46. Lightman, S. M., Utley, A. & Lee, K. P. Survival of long-lived plasma cells (LLPC): Piecing together the puzzle. Front. Immunol. 10, 1–12 (2019).

47. Bonsignori, M. et al. HIV-1 envelope induces memory B cell responses that correlate with plasma antibody levels after envelope gp120 protein vaccination or HIV-1 infection. J. Immunol. 183, 2708–2717 (2009).

48. Landais, E. et al. Broadly Neutralizing Antibody Responses in a Large Longitudinal Sub-Saharan HIV Primary Infection Cohort. PLoS Pathog. 12, 1–22 (2016).

49. Remmel, J. L. & Ackerman, M. E. Rationalizing Random Walks: Replicating Protective Antibody Trajectories. Trends Immunol. 42, 186–197 (2021).

50. Steichen, J. M. et al. A generalized HIV vaccine design strategy for priming of broadly neutralizing antibody responses. Science 366, (2019).

51. Ingale, J. et al. High-density array of well-ordered HIV-1 spikes on synthetic liposomal nanoparticles efficiently activate B cells. Cell Rep. 15, 1986–1999 (2016).

52. Martinez-Murillo, P. et al. Particulate array of well-ordered HIV clade C Env trimers elicits neutralizing antibodies that display a unique V2 cap approach. Immunity 46, 804–817 (2017).

53. Stano, A. et al. Dense Array of Spikes on HIV-1 Virion Particles. J. Virol. 91, 1–19 (2017).

54. Gonelli, C. A. et al. Immunogenicity of HIV-1-based virus-like particles with increased incorporation and stability of membrane-bound env. Vaccines 9, 1–36 (2021).

55. Mohsen, M. O., Zha, L., Cabral-Miranda, G. & Bachmann, M. F. Major findings and recent advances in virus–like particle (VLP)-based vaccines. Semin. Immunol. 34, 123–132 (2017).

56. Pan, Y.-W., Scarlett, J. M., Luoh, T. T. & Kurre, P. Prolonged Adherence of Human Immunodeficiency Virus-Derived Vector Particles to Hematopoietic Target Cells Leads to Secondary Transduction In Vitro and In Vivo. J. Virol. 81, 639–649 (2007).

57. Kuate, S. et al. Immunogenicity and efficacy of immunodeficiency virus-like particles pseudotyped with the G protein of vesicular stomatitis virus. Virology 351, 133–144 (2006).

58. Michelini, Z. et al. Development and use of SIV-based Intgrase defective lentiviral vector for immunization. Vaccine 23, 4622–4629 (2009).

59. Trobridge, G. D. et al. Cocal-pseudotyped lentiviral vectors resist inactivation by human serum and efficiently transduce primate hematopoietic repopulating cells. Mol. Ther. 18, 725–733 (2010).

60. Finkelshtein, D., Werman, A., Novick, D., Barak, S. & Rubinstein, M. LDL receptor and its family members serve as the cellular receptors for vesicular stomatitis virus. Proc. Natl. Acad. Sci. U. S. A. 110, 7306–7311 (2013).

61. Berger, G. et al. A simple, versatile and efficient method to genetically modify human monocyte-derived dendritic cells with HIV-1-derived lentiviral vectors. Nat. Protoc. 6, 806–816 (2011).

62. Montefiori, D. C. Evaluating neutralizing antibodies against HIV, SIV, and SHIV in luciferase reporter gene assays. Curr. Protoc. Immunol. Chapter 12, Unit 12.11 (2005).

63. Montefiori, D. C. Measuring HIV neutralization in a luciferase reporter gene assay. Methods Mol. Biol. 485, 395–405 (2009).

64. Bentler, P. M. & Weeks, D. G. Linear structural equations with latent variables. Psychometrika 45, 289–308 (1980).

65. STATA. Structural Equation Modelling Reference Manual. Release 16. College Station, TX: Stata Press (2019).

