## Supplemental Material for "Persistent Immunogenicity of Integrase Defective Lentiviral Vectors delivering membrane tethered Native-Like HIV-1 Envelope Trimers"

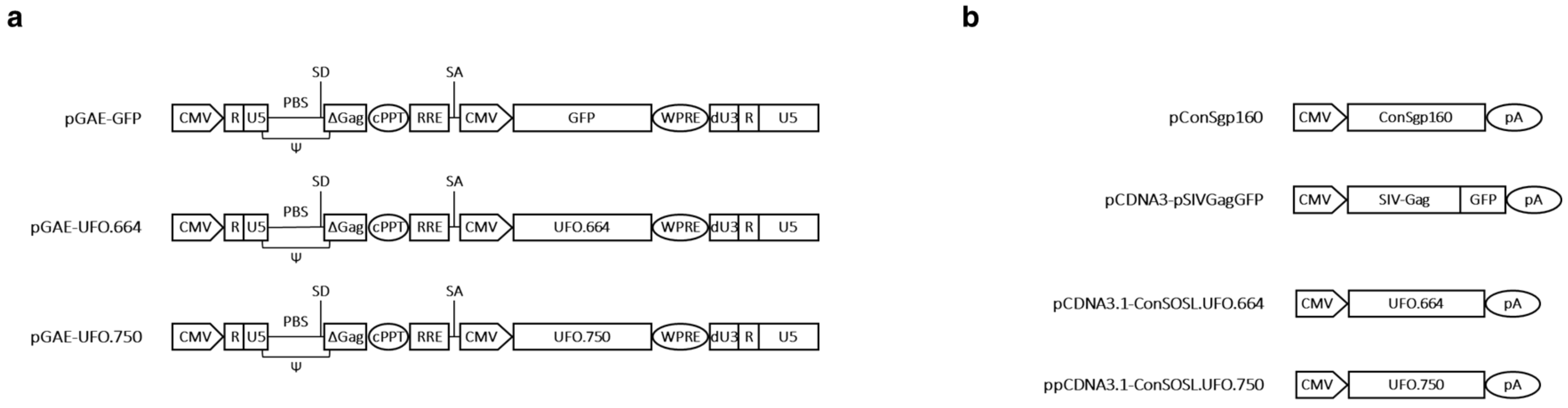


**Supplementary Figure 1. Schematic representation of plasmids used in this study.** (**a**) SIV-based lentiviral transfer vectors expressing GFP, ConSOSL.UFO.664 or ConSOSL.UFO.750. (**b**) pcDNA3 plasmids expressing ConSgp160, SIV-Gag fused to GFP, ConSOSL.UFO.664 or ConSOSL.UFO.750. CMV, cytomegalovirus immediate-early promoter; R, repeat element; U5, 5' untranslated region; U3, 3' untranslated region; PBS, primer binding site; SD, splice donor site; Ψ, packaging signal; cPPT, central polypurine tract; RRE, Rev response element; SA, splice acceptor site; dU3, SIN deletion in U3 region of 3' LTR; WPRE, woodchuck hepatitis virus post-transcriptional regulatory element. See Methods for details on construction.

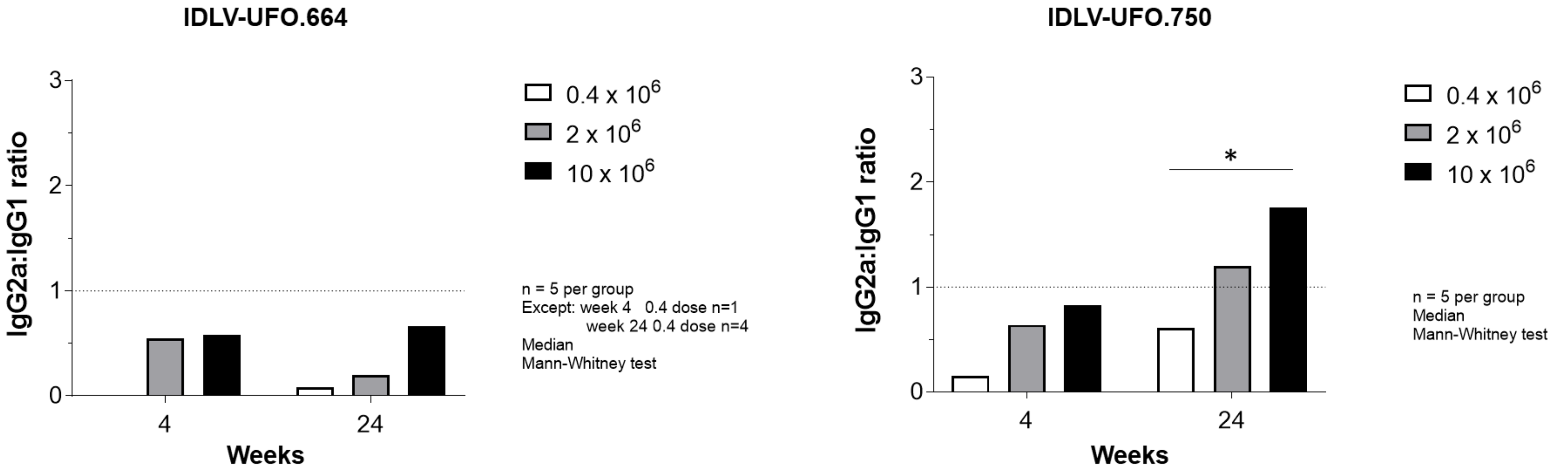


**Supplementary Figure 2. IDLV-UFO.750 induced a more pronounced Th1 response than IDLV-UFO.664 in immunized mice.** Sera from BALB/c mice vaccinated with escalating doses of IDLV-UFO.664 or IDLV-UFO.750 were analyzed by ELISA for IgG1 and IgG2a ConSOSL.UFO-specific responses at weeks 4 and 24 from immunization. Results are shown as IgG2a/IgG1 ratio. *$p<0.05$, Mann-Whitney test.

**Table S1.** Anti-ConSOSL.UFO.664 IgG in mucosal secretions from immunized monkeys.

| | Animal ID | AS304 | | AT777 | | AU018 | | AU955 | | AU989 | |
|---|---|---|---|---|---|---|---|---|---|---|---|
| Vaccine regimen | Week | Saliva | Rectal | Saliva | Rectal | Saliva | Rectal | Saliva | Rectal | Saliva | Rectal |
| IDLV-UFO750 | 0 | 0,02 | 0 | 0,01 | 0 | 0 | 0 | 0 | 0 | 0 | 0 |
| | 2 | 0,11 | 0 | 0,09 | 0,04 | 0.01 | 0 | 0 | 0 | 0 | 0 |
| | 6 | 0 | 0 | 0 | 0 | 0 | 0 | 0 | 0 | 0 | 0 |
| | 10 | 0 | 0 | 0 | 0 | 0 | 0 | 0 | 0 | 0 | 0 |
| | 14 | 0 | 0 | 0 | 0 | 0 | 0 | 0 | 0 | 0 | 0 |
| | 18 | 0 | 0 | 0,03 | 0 | 0 | 0 | 0 | 0 | 0 | 0 |
| | 22 | 0 | 0 | 0 | 0 | 0,13 | 0 | 0 | 0 | 0 | 0 |
| | 27 | 0 | 0 | 0 | 0 | 0 | 0 | 0 | 0 | 0 | 0,01 |
| | 33 | 0 | 0 | 0 | 0 | 0 | 0 | 0 | 0 | 0,13 | 0,02 |
| IDLV-UFO750 | 37 | 0 | 0 | 0 | 0 | 0 | 0 | 0 | 0 | 0 | 0 |
| | 39 | 0,19 | 0 | 0,32 | 0 | 0 | 0 | 0,75 | 0 | 0,68 | 0,09 |
| | 43 | 0,17 | 0 | 0,27 | 0 | 0 | 0 | 0,16 | 0 | 0,43 | 0,03 |
| | 47 | 0 | 0 | 0,09 | 0 | 0 | 0 | 0 | 0,06 | 0 | 0,02 |
| | 52 | 0 | 0 | 0 | 0 | 0 | 0 | 0 | 0 | 0 | 0 |
| | 57 | 0 | 0 | 0 | 0 | 0 | 0 | 0 | 0 | 0 | 0 |
| ConM SOSIP.v7 + MPLA | 62 | 0 | 0 | 0 | 0 | 0 | 0 | 0 | 0 | 0,08 | 0 |
| | 64 | 0,17 | 0,02 | 0,16 | 0 | 0 | 0 | 0,98 | 0 | 0,34 | 0,01 |
| | 69 | 0,07 | 0 | 0,04 | 0 | 0 | 0 | 0 | 0,02 | 0 | 0,06 |
| ConM SOSIP.v7 + MPLA | 74 | 0 | 0 | 0 | 0 | 0 | 0 | 0,05 | 0,24 | 0 | 0 |
| | 76 | 0,15 | 0 | 0,18 | 0 | 0 | 0 | 1,43 | 0 | 0,38 | 0,05 |
| | 80 | 0,05 | 0 | 0,02 | 0 | 0 | 0 | 0,21 | 0 | 0,10 | 0 |
| | 85 | 0 | 0 | 0 | 0 | 0 | 0 | 0 | 0 | 0 | 0 |
| | 89 | 0 | 0 | 0,07 | 0 | 0 | 0 | 0 | 0 | 0 | 0 |
| | 94 | 0 | 0 | 0 | 0 | 0 | 0 | 0 | 0 | 0 | 0 |

Saliva and rectal swabs from immunized monkeys were collected at the indicated time points and analyzed by capture ELISA for the presence of anti-ConSOSL.UFO specific Abs. Results are expressed as the % of specific anti-ConSOSL.UFO.664 IgG compared to total IgG.
